# Maltose aids in the growth of *Salmonella* Typhimurium in nitrogen-deficient conditions

**DOI:** 10.64898/2026.01.15.699617

**Authors:** Kirti Parmar, Yogyta Kumari, Muralidhar Nayak M, Kapudeep Karmakar, Sumanta Bagchi, Dipshikha Chakravortty

## Abstract

Enteric pathogens, such as *Salmonella,* encounter uncertain nutrients in both the host and the environment. Nutrients such as carbon and nitrogen are essential for growth, and when these nutrients are limited, bacteria have developed various mechanisms to overcome stress and thrive in nutrient-rich environments. In this study, we investigated the role of the disaccharide sugar maltose in the growth and pathogenicity of *Salmonella* Typhimurium (STM) under nitrogen-deficient culture conditions. Throughout the study, we used Burk’s medium to mimic nitrogen-deficient conditions. STM growth in Burk’s medium was minimal, and maltose supplementation enhanced STM growth in both liquid and solid Burk’s media. The STM grown in Burk’s medium had a reduced bacterial cell length, which increased upon the addition of maltose and became comparable to that of LB-grown STM WT. Growth was dependent on the NtrBC two-component system and sigma factor RpoE. Deletion of *ntrB* and *rpoE* significantly reduced the growth in Burk’s medium during the log phase. *Salmonella* grown in Burk’s medium with maltose had reduced invasion in Caco-2 cells, phagocytosis in RAW 264.7 macrophages, and organ burden in C57BL/6 mice. Our findings revealed the role of maltose in enhancing the growth and length of *Salmonella* under nitrogen-deficient conditions. However, *Salmonella* grown in Burk’s medium with maltose is defective in pathogenesis.

## 1. Introduction

Bacterial growth often occurs in environments with limited nutrient availability. Various bacteria use different physiological adaptations to overcome this nutrient limitation. Enhanced expression of siderophores to scavenge the limited nutrients such as iron, spore formation in Gram-positive bacteria, growth arrest and stringent response in *E.coli,* increased length and appendages in *Caulobacter crescentus* to enhance nutrient uptake(Estrela *et al*., 2025).

Nitrogen and carbon are the essential nutrients required for growth. Bacteria respond to nitrogen deficiency by remodelling gene expression. Nitrogen stress is sensed by the proteins GlnB and GlnK of the PII signalling family, leading to the activation of the nitrogen regulatory protein C (NtrC), which increases the expression of nitrogen transporters in *E.coli,* resulting in nitrogen influx (Switzer *et al*., 2018). NtrC also binds to the promoter of the eukaryote-like serine-threonine kinase *yeaG* in *E.coli,* which controls population heterogeneity and deletion, leading to compromised survival during prolonged nitrogen starvation owing to its influence on the general stress response regulator RpoS (Figueira *et al*., 2015). NtrC also activates *relA,* which couples the nitrogen stress response with the stringent response. A stringent response leads to metabolically inactive cells or the formation of persister cells, resulting in increased resistance to antibiotics such as ciprofloxacin (Brown *et al*., 2014, Sanchuki *et al*., 2017, Brown, 2019). NtrC is also needed for the long-term survival (150 days) of *Salmonella* in nitrogen-deficient conditions (Mishra & Shashidhar, 2022).

In addition to the above reports, *E.coli* growth on poor or rare sources of nitrogen, such as amino acids, exhibits diauxic growth when grown on two different nitrogen sources (Atkinson *et al*., 2002). Additionally, there is a preference for sugars such as maltose or maltodextrins over glucose for growth in poor nitrogen sources (Bren *et al*., 2016). Glucose uptake is inhibited in nitrogen-starved *E.coli* due to the inhibition of the PtsI protein by alpha-ketoglutarate, and overexpression of PtsI leads to glucose uptake (Chubukov *et al*., 2017). Similarly, in actinomycetes, there is an increased utilisation and uptake of non-phosphotransferase system carbon sources, such as maltose, under nitrogen-deficient conditions (Liao *et al*., 2015).

All of the above studies demonstrated growth arrest and metabolically inactive cells during nitrogen starvation when using minimal media with glucose as the carbon source, as well as some studies on the preference for the sugar maltose. In our study, we used a nitrogen-free Burk’s medium to isolate *Azotobacter* and observed the growth of *Salmonella* Typhimurium when supplemented with maltose. We attempted to understand the survival of *Salmonella* during nitrogen starvation, as nutrient scarcity is an effective defence mechanism of the host and is also present in the environment outside the host. Overcoming stress may be one reason why *Salmonella* is a successful pathogen.

## 2. Materials and Methods-

### 2.1 Bacteria and growth conditions

Wild-type *Salmonella* Typhimurium 14028s (STM WT) was obtained from Professor Michael Hensel (Universität Osnabrück). STM WT was revived from glycerol stock stored at -80°C and streaked on LB agar; the respective antibiotic-containing plates were used for the knockout strains. All the bacterial strains were grown in LB supplemented with the required antibiotics at 37°C and 170 rpm overnight. The strain containing pkD46 was grown at 30°C and 170 rpm.

To mimic the nitrogen-deficient state, Burk’s medium (Hi-media, catalogue number M707), supplemented with various carbon sources and ammonium chloride, was used. Burk’s medium was composed of magnesium sulphate 0.200, dipotassium hydrogen phosphate 0.800, potassium dihydrogen phosphate 0.200, calcium sulphate 0.130, iron (III) chloride 0.00145, sodium molybdate 0.000253, and sucrose 20.000. All the components are in grams per litre.

Overnight cultures of different bacterial strains were pelleted, washed with 1X PBS, and resuspended in 1X PBS. The bacterial culture was subcultured at a 1:1000 ratio under various conditions, as described in the figure. The starting optical density (OD) of the growth curves was observed to be 0.05. The growth was measured over 12 hours at 3-hour intervals using optical density at 595nm at 37°C and 170 rpm. Samples were transferred to 96-well plates in 3 replicates, and the absorbance was measured using a Tecan instrument.

The growth of various strains was measured using Bioscreen at a sub-culture ratio of 1:100, and was performed at 1-hour intervals for 24 hours. The starting OD of the cultures was observed to be 0.15-0.2. A list of the bacterial strains used in this study is shown in Table 1.

**Table 1.**
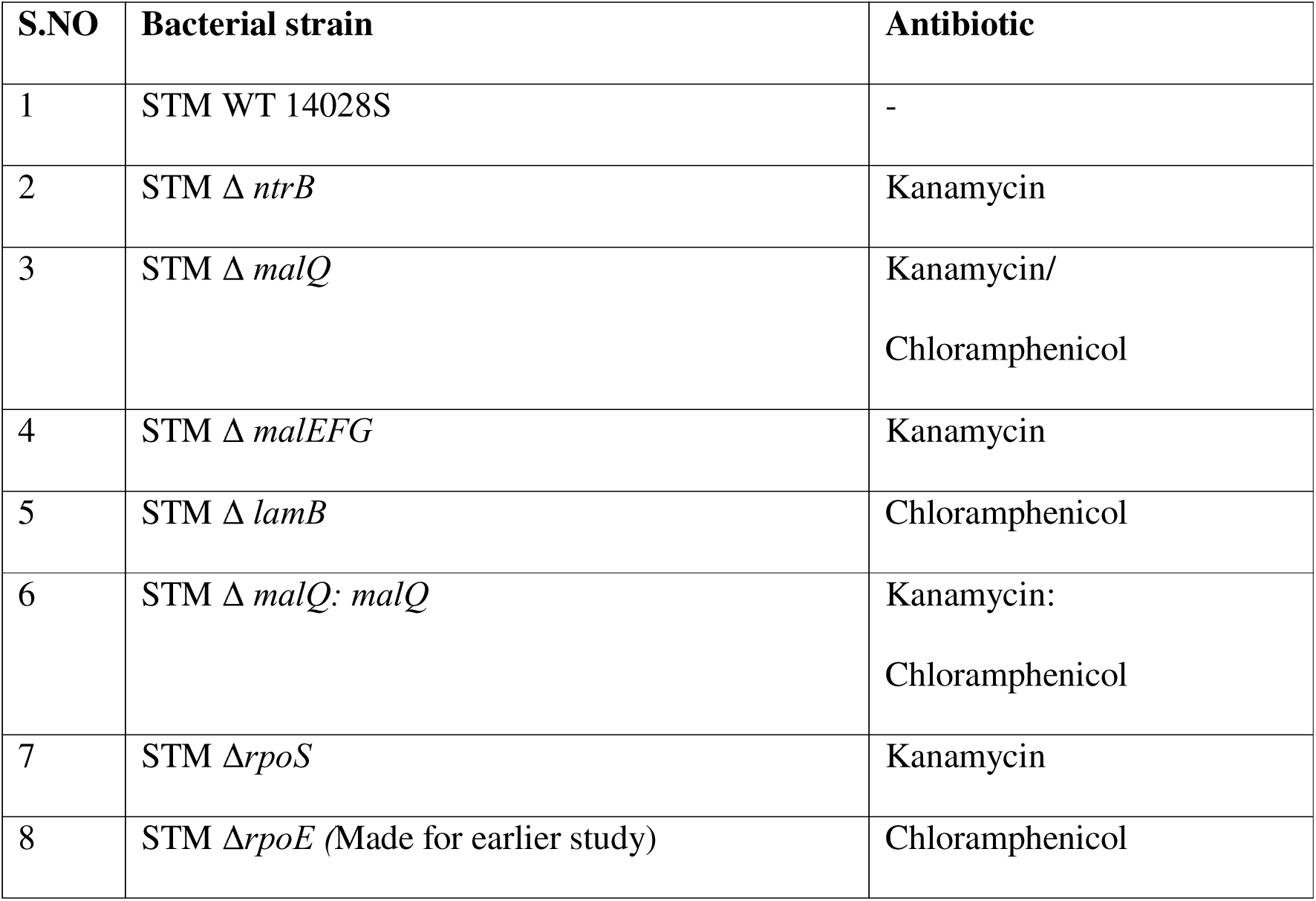
List of bacterial strains used in the study.

### 2.2 Growth on solid media

Petri plates were prepared with Burk’s medium (B), Burk’s medium with 0.2% maltose (BM), Burk’s medium with 0.2% maltose and 10 mM ammonium chloride (BMN), all supplemented with 1.5% agar. Maltose was added after autoclaving the medium before pouring the plates. An overnight culture of STM WT in LB media was pelleted, washed with 1X PBS, and resuspended in 1X PBS. Bacterial cultures were diluted and plated onto petri plates. Plates were incubated for 24 hours, and images were acquired using Chemidoc.

### 2.3 Elemental analysis

To check the nitrogen status, elemental analysis was performed in LB, Burk’s medium and maltose to confirm the presence of any nitrogen contamination using a combustion based CHNS analyser at 650°C. Nitrogen concentration was analysed using combustion gas oxygen (<99.99%) and carrier gas helium (<99.99%) using a standard against sulphenamide. Elemental analysis was performed by gas chromatography (Vario EL Cube, Elementar, Germany). The percentage obtained for elemental nitrogen was weight by weight.

### 2.4 Scanning electron microscopy

STM WT was grown overnight in LB media and pelleted down, washed with 1X PBS, and resuspended in 1X PBS. WT was subcultured in different media conditions at a 1:1000 ratio (5 µL in 5 mL) and incubated for 6 hours to obtain a log-phase culture. The log phase culture was pelleted and resuspended in 1X PBS. Subsequently, 50 µL of the sample was added to a coverslip and air-dried. The dried samples were fixed with 3.5 % glutaraldehyde overnight at 4°C. The next day, samples were washed with Milli-Q and treated with a gradient of various alcohol concentrations (10%, 30%, 50%, 70%, 90%, and 100%) prepared in Milli-Q for 2-3 minutes to achieve dehydration. The samples were desiccated for 30 minutes, mounted on copper stubs using carbon tape, and then gold-coated. Images were acquired using an ESEM Quanta microscope, and the lengths of the bacteria were analysed using ImageJ software.

### 2.5 Construction of knockout strains

Various genes were knocked out using a one-step chromosomal gene inactivation protocol described by Datsenko and Wanner (Datsenko & Wanner, 2000). The knockout primers were designed to contain a 40-bp homologous sequence to the gene to be knocked out, along with a 20-bp extension for the antibiotic cassette. The gene encoding for the antibiotic was amplified from *E.coli* using the pKD3 (chloramphenicol) or pKD4 (kanamycin) plasmid. The PCR product was purified using the chloroform-isopropanol method and then analysed on an agarose gel to verify its size.

For electroporation, STM pKD46 bacterial cells (which contain a lambda red recombinase system under an arabinose-inducible promoter) were grown to the log phase in the presence of arabinose and ampicillin, and the absorbance was adjusted to 0.3. The cells were then made electrocompetent by washing three times with MilliQ water and three times with 10% glycerol at 4°C. The PCR product was incubated with electrocompetent cells for 30 minutes, followed by electroporation. Immediately after electroporation, 1 mL of LB medium was added to the cells for recovery and maintained at 37°C for 1 hour. The bacteria were pelleted and plated onto antibiotic-containing plates. Gene knockout was confirmed by patching the bacteria onto a fresh plate containing antibiotics and performing PCR on the resulting colonies. Primers used in this study are listed in Supplementary Table 1.

### 2.6 Eukaryotic cell lines and growth conditions

Cell line Caco-2 used in this study was maintained in Dulbecco’s Modified Eagle’s Medium (DMEM, Sigma-Aldrich) supplemented with 10% FBS (fetal bovine serum, Gibco), 1% sodium pyruvate, 1% penicillin-streptomycin, and 1% non-essential amino acids at 37°C in the presence of 5% CO_2_. Upon confluency, cells were harvested by trypsin-EDTA treatment, counted using a haemocytometer, and then seeded in a 24-well plate.

RAW 264.7 macrophages were maintained in DMEM supplemented with 10% FBS and 1% penicillin-streptomycin. Cells were grown to confluence, harvested with a cell scraper, and counted with a haemocytometer. An equal number of cells were seeded in 24-well plates.

### 2.7 Invasion assay

One lakh seeded Caco2 cells were infected with bacteria at an MOI of 10, and a log phase culture was used to infect the epithelial cells (Caco2 cells). Cells were centrifuged at 800 rpm for 5 minutes and then incubated at 37°C with 5% CO2 for 25 minutes. Cells were then washed with 1X PBS to remove all extracellular bacteria, and media containing 100µg/ml gentamicin in DMEM with FBS was added for 1 hour. Similarly, 25 µg/ml of gentamicin in DMEM media with FBS was added, and the cells were incubated until lysis. Cells were lysed with 0.1% Triton X-100 at 2 hours. Bacteria were plated on the LB Agar, incubated at 37°C for 12 hours, and colonies were counted. To calculate the percentage invasion, bacterial CFU at 2 hours were divided by the pre-inoculum CFU.

### 2.8 Phagocytosis assay

One lakh seeded RAW 264.7 cells were infected with bacteria at an MOI of 10, and a stationary phase was used for infection. A protocol similar to the one described above was followed. To calculate the percentage of phagocytosis, bacterial CFU at 2 hours were divided by the pre-inoculum CFU.

### 2.9 Proliferation assay

One lakh seeded Caco2 cells/ RAW 264.7 cells were infected with bacteria at an MOI of 10 as mentioned above. Cells were centrifuged at 800 rpm for 5 minutes and then incubated at 37°C with 5% CO2 for 25 minutes. Cells were then washed with 1X PBS to remove all extracellular bacteria, and media containing 100µg/ml gentamicin in DMEM with FBS was added for 1 hour. Similarly, 25 µg/ml of gentamicin in DMEM media with FBS was added, and the cells were incubated until lysis. Cells were lysed with 0.1% Triton X-100 at 2 hours and at a later time point of 16 hours. Bacteria were plated on the LB Agar, incubated at 37°C for 12 hours, and colonies were counted. To calculate the fold proliferation, the bacterial CFU at 16 hours was divided by the CFU at 2 hours.

### 2.10 Alpha-ketoglutarate estimations

A protocol from Rabinowitz et al. was followed (Rabinowitz & Kimball, 2007). Briefly, STM WT was grown in LB medium overnight and centrifuged at 6000 rpm for 5 min to pellet the bacteria. Then, the cells were washed with 1x PBS and resuspended in 1x PBS. STM WT was subcultured at a 1:100 ratio in different conditions (Burk’s medium, Burk’s medium + 0.2% maltose, Burk’s medium + 0.2% maltose + 10mM ammonium chloride, LB medium) for 5 hours. Log-phase culture was harvested at 5,000 rpm for 5 minutes. The pellet was resuspended in 300 μL of extraction solvent. Acetonitrile extraction was performed by adding acetonitrile, water, and formic acid (0.1M) containing the extraction solvent at an 80:20 ratio. Samples were incubated at 4°C for 15 minutes and then centrifuged at 13000 rpm, 4°C to separate insoluble materials from the extracted metabolites. The alpha-ketoglutaric acid standard was prepared by dissolving it in Milli Q water. Standard solutions of various concentrations were prepared in acetonitrile and water (70:30). All the samples were analysed using mass spectroscopy, and the concentration of alpha-ketoglutarate was determined. The obtained values were then normalised to the CFU of the bacteria in the sample.

UPLC Conditions and ESI-Triple quad Mass Spectrometry

The sample extracts were analysed using a UPLC–ESI–MS/MS system (UPLC, SHIMADZU, LCMS-8045) with Lab Solution software. The UPLC conditions are described as follows: Mobile phase A: water containing 0.1% formic acid, mobile phase B: acetonitrile (MS grade, Honeywell, Cat. No: No:15040-2.5L) with an isocratic elution of 70 % mobile phase A and 30% mobile phase B with a Flow rate of 0.3 mL/ min. The injection volume was 40 µL for samples and 10 µL for standards, with a total run time of 10 minutes. The column temperature was 25°C, and the autosampler temperature was 15°C. The ESI source parameters were set: ion spray voltage 4000 V (negative ion mode); other set parameters were interface temperature 300 °C, desolvation temperature 526 °C, nebulizing gas (nitrogen) flow rate 3 L/min, Dry gas (nitrogen) flow rate 10L/min, heating gas (nitrogen) flow rate of 10L/min, heat block temperature 400 °C, CID gas (Argan gas 99.9995% for collision) flow limit set to 230kPa. The ion reaction monitoring (MRM) mode was then selected using mass spectrometry scanning, and the specific mass-to-charge ratio (m/z) of alpha-ketoglutaric acid (145.05 m/z) was set using QQQ scans in MRM mode (145.05 > 101.00). The precursor ion, 145.05 m/z and product ion, 101.00 m/z, were selected for ion section transfer using a based peak intensity (BPI) scan. Operational parameters, such as the ion source gas, scan time, and collision energy, were set to achieve optimal signal intensity and peak separation. The time of each scan event was set to 0.309 sec.

### 2.11 RNA isolation and cDNA Synthesis

STM WT was grown overnight in LB medium and washed with 1x PBS before subculturing (1:100) in different media (Burk’s medium + 0.2% maltose, Burk’s medium + 0.2% maltose + 10mM ammonium chloride, LB medium). A log phase culture was collected after 6 hours to analyse the expression of various invasion genes. The bacterial culture was pelleted by centrifugation at 6000 rpm for 10 minutes, and 1 ml of TRIzol (Takara) was added to the pellet. The pellet was resuspended in TRIzol, and the samples were stored at -80°C until processing. RNA was extracted using chloroform, centrifuged at 12,000 rpm for 15 minutes, and then precipitated by adding an equal volume of isopropanol. The supernatant was then incubated at room temperature for 30 minutes, followed by centrifugation at 12,000 rpm for 30 minutes. The obtained pellet was washed with 70% ethanol in DEPC water and air-dried. The RNA was dissolved in DEPC water. The quality and quantity of RNA were checked using Nanodrop and 2% agarose gel electrophoresis. DNase treatment (Thermo Fisher Scientific) was performed on the isolated RNA at 37°C for 1 hour, followed by heat inactivation at 65°C for 15 minutes. DNase-treated RNA was converted to cDNA using a PrimeScript Reagent RT kit (Takara). qRT-PCR was performed using a TB Green RT-qPCR Kit in a real-time PCR system. For the analysis of RT data, 16S rRNA was used as the reference gene to calculate the delta Ct (ΔCt) of the samples. ddct was calculated by comparing the two different samples.

### 2.12 Animal experiments-

Five-to six-week-old C57BL/6 male mice were used for survival and organ burden determination. Mice were orally gavaged with 10^8^ (100 µL) bacteria, and their weight and health were monitored to calculate their weight reduction percentage and survival. To determine the organ burden in mice, mice were orally gavaged with 10^7^(100 µL) bacteria, euthanised, and various organs, including blood, liver, and mesenteric lymph node (MLN), were harvested. Blood was isolated via a heart puncture. Mouse organs were homogenised and plated on *Salmonella Shigella* agar to determine the bacterial burden in various organs.

### 2.13 Statistics

The Statistical tests used in the experiments are mentioned in the figure legends. The Mann-Whitney test was performed for animal experiments. All the data were plotted and analysed using GraphPad Prism 8.

## 3. Results

### 3.1 Maltose in nitrogen-deficient media enhances the growth of *Salmonella*

The Burk medium, commonly used to isolate diazotrophs, was employed to assess the growth of *Salmonella* under nitrogen-deficient conditions (Park *et al*., 2005). Since low concentrations of contaminating nitrogen may be present in every medium, we performed elemental analysis of Burk’s medium, maltose, and LB media. It was observed that maltose had no traceable amounts of nitrogen, whereas 0.2 - 0.4% (w/w) nitrogen was found in Burk’s media compared to 8 – 10% (w/w) in LB media **(Supplementary Figure 1A).** The nitrogen concentration in the Burk’s media translates to 4.5 mM. Spectroscopic analysis of bacterial growth was performed using optical density in Burk’s medium, supplemented with varying concentrations of glucose and maltose. Growth of *Salmonella* in nitrogen-deficient Burk’s medium was poor but significantly enhanced when supplemented with 0.2% maltose **(Supplementary Figure 1B).** Thus, it was concluded that supplementation with 0.2% maltose enhances the growth of *Salmonella* under nitrogen-deficient conditions. In the literature, 10 mM ammonium chloride is defined as a nitrogen-rich condition that mimics a nitrogen-rich condition (Schumacher *et al*., 2013). Upon performing a growth analysis at 1:100 and 1:1000 subculture ratios, we observed the highest growth of *Salmonella* in nutrient-rich LB medium, as expected, and weak growth in Burk’s medium. There was increased growth of *Salmonella* in maltose supplemented Burk’s medium, but there was no significant growth advantage when nitrogen was added to Burk’s medium with maltose **(Figure 1A).** Similarly, when STM was grown in solid media, there was a reduced colony size on Burk’s medium, which increased with the supplementation of maltose alone or in combination with ammonium chloride **(Figure 1B).** Furthermore, to validate that the increased growth of *Salmonella* in Burk when supplemented with maltose was due to maltose uptake and metabolism, the maltose metabolism gene (*malQ)* and maltose transporters (*malEFG*-ABC transporter, *lamB -* maltoporin) were deleted. Analysis of the growth of these strains in Burk’s medium supplemented with maltose (BM) showed that STM Δ*malQ* and STM Δ*malEFG* exhibited a significant reduction in growth **(Figure 1C, 1D).**

**Figure 1.**
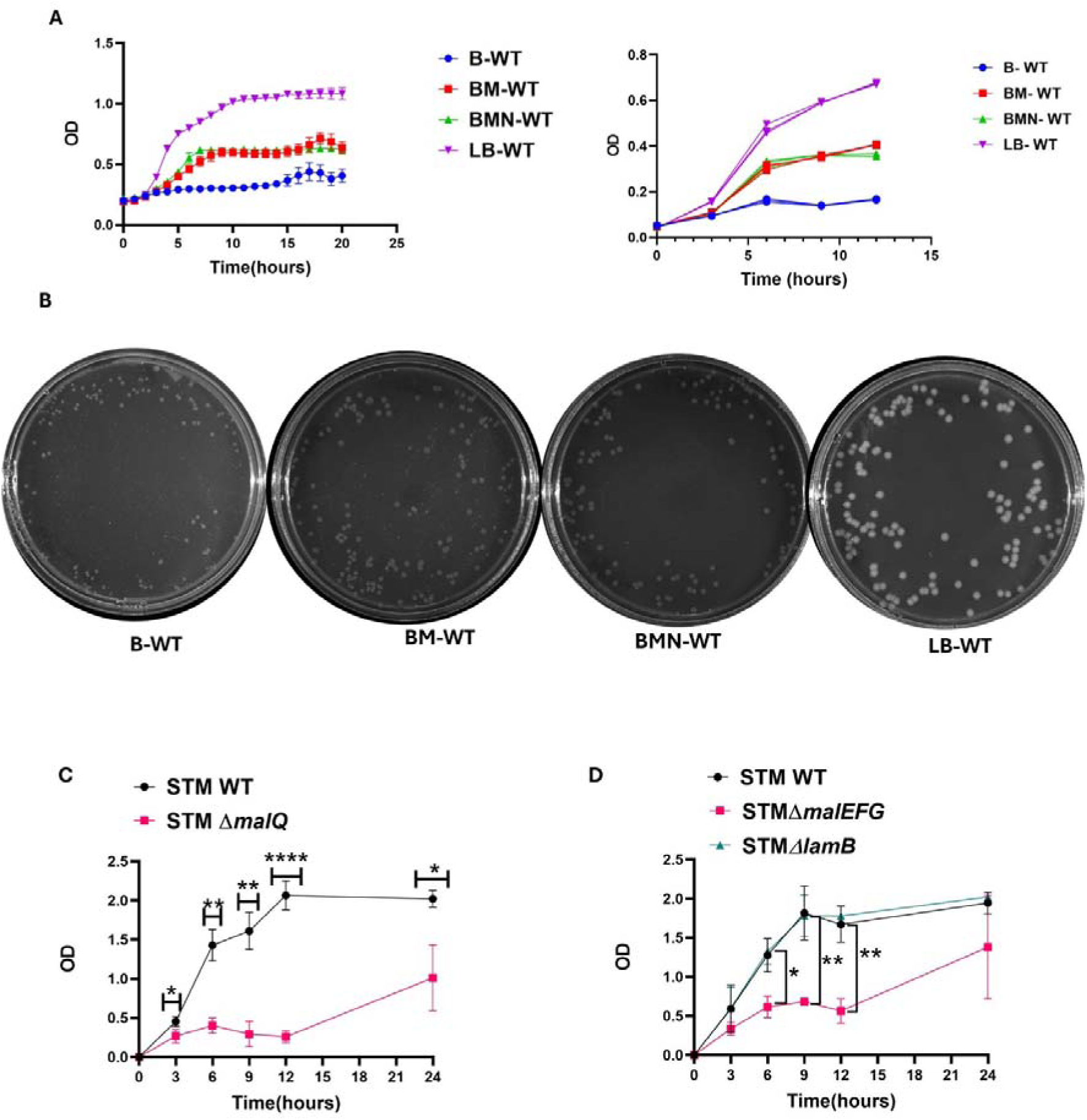
*Salmonella* growth in a nitrogen-deficient medium was enhanced by maltose supplementation. A. Growth curve analysis by absorbance of STM WT grown under four different conditions at various time points in a subculture ratio of 1:100 (left) and 1:1000(right). Media conditions-B (Burk’s medium), BM (Burk’s medium + 0.2% maltose), BMN (Burk’s medium + 0.2% maltose +10mM ammonium chloride), LB (Luria Bertani medium). Data are representative of N=3,n=3, presented as mean±SD. B. Growth of STM WT on different agar media plates from the same culture. Data are representative of N = 3, n = 2. C. Growth curve analysis by absorbance at various time points of STM WT and STM Δ*malQ* at a subculture ratio of 1:100. Unpaired Student’s t-test was used for the analysis. Data are representative of N=3, n=2, presented as mean±SD. D. Growth curve analysis by absorbance at various time points for STM WT, STM Δ*lamB* and STM Δ*malEFG* at a subculture ratio of 1:100. Unpaired Student’s t-test was used for the analysis. Data are representative of N=3, n=2, presented as mean±SD.

Bacterial cell size is dependent on nutrient availability, and nutrient limitation leads to a decrease in bacterial length (*Salmonella*) and diameter (*Bacillus Subtilis* and *Staphylococcus aureus*) (Watson *et al*., 1998, Chien *et al*., 2012). In contrast, in *Bacteroides thetaiotaomicron*, there is an increase in size under sugar limitation (Rangarajan *et al*., 2020). Next, we determined the length of the bacteria grown in the above conditions, STM grown in Burk’s medium (1.032 µm) was shorter compared to bacteria grown in Burk’s medium with supplementation of maltose alone (BM-1.44 µm) or in combination with ammonium chloride (BMN-1.35 µm) and LB media (1.37 µm) (**Figure 2A, B**). It was previously reported that nitrogen deficiency in bacteria is associated with enrichment of the metabolite alpha-ketoglutarate and a reduction in glutamine (18), as determined by analysis of alpha-ketoglutarate levels in bacteria grown under these conditions. It was observed that with increased growth of *Salmonella* in Burk’s medium supplemented with maltose (BM), there were higher alpha-ketoglutarate levels in bacteria grown in the Burk’s medium (B), Burk’s medium supplemented with maltose, ammonium chloride (BMN), and LB media **(Figure 2C).** In conclusion, maltose supplementation facilitates *Salmonella* growth under nitrogen-deficient conditions, enhances bacterial length, and leads to higher alpha-ketoglutarate levels.

**Figure 2.**
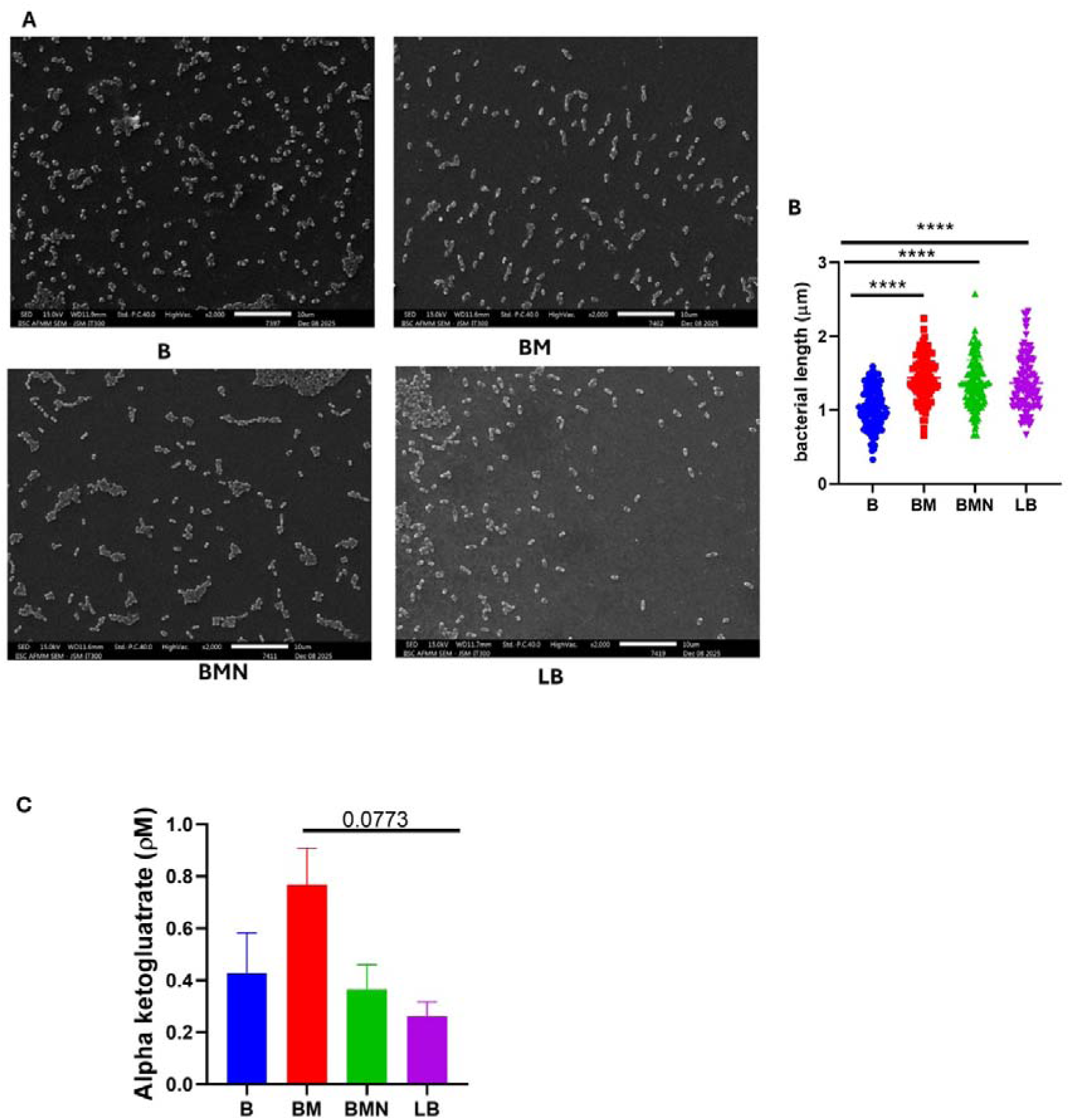
STM length and alpha-ketoglutarate levels when grown in a nitrogen-deficient medium. A. Representative SEM images of STM WT grown until log phase under different conditions at a subculture ratio of - B (Burk’s medium), BM (Burk’s medium + 0.2% maltose), BMN (Burk’s medium + 0.2% maltose +10mM ammonium chloride), LB (Luria Bertini medium).N=2 B. Quantification of bacterial length in A: One-way ANOVA was used for analysis. Data are presented as the mean ± SD, N = 2. Approximately 110 bacterial cells were counted for each condition. C. Estimation of alpha-ketoglutarate levels in STM WT grown under four conditions mentioned above in the log phase. Statistical analysis was performed by using an unpaired Student’s t-test. N=2, presented as the mean±SEM.

### 3.2 *Salmonella* growth in a nitrogen-deficient condition is dependent on *ntrB* and rpoE

Nitrogen deficiency, accompanied by an increase in alpha-ketoglutarate, triggers the activation of the NtrBC two-component system, which in turn activates other nitrogen-associated genes (Schumacher *et al*., 2013). We quantified the transcript levels of *ntrB* in Burk’s medium supplemented with nitrogen and maltose (BMN) compared to Burk’s medium supplemented with maltose (BM) and found *ntrB* was downregulated at all time points from 3-12 hours **(Figure 3A)**. Nitrogen assimilation from ammonia in bacteria is catalysed by two important enzymes, glutamine synthetase, encoded by *glnA*, and glutamate dehydrogenase, encoded by *gldH (Yan, 2007). glnA* is activated during nitrogen scarcity, and glutamate dehydrogenase is active when nitrogen concentration is high (Atkinson *et al*., 2002, Kumar & Shimizu, 2010). The mRNA levels of *gldH* were downregulated in Burk’s medium supplemented with nitrogen and maltose (BMN) compared to Burk’s medium supplemented with maltose (BM), but remained constant throughout 3-12 hours **(Figure 3B).** Analysis of the mRNA of *glnA* in Burk’s medium supplemented with nitrogen and maltose (BMN) showed reduced expression at later time points of 6, 9, and 12 hours compared to Burk’s medium supplemented with maltose (BM), suggesting that nitrogen is not limiting on the addition of ammonium chloride **(Figure 3C).** The downregulated expression of genes *ntrB, glnA* and *gldH* showed that the addition of 10mM ammonium chloride led to nitrogen-rich conditions in Burk’s medium supplemented with maltose.

**Figure 3.**
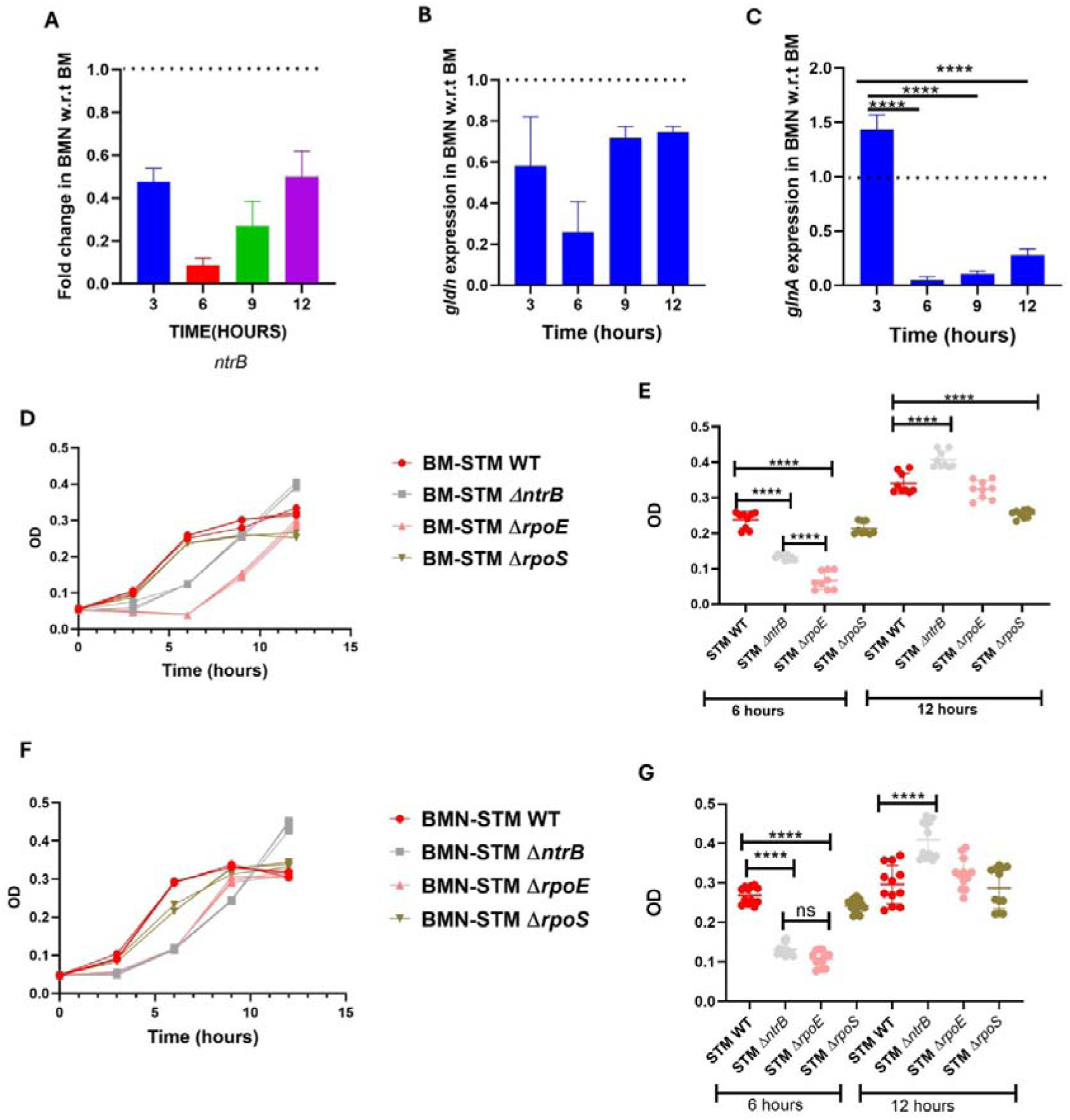
Determining the factors responsible for *Salmonella* growth. A-C. mRNA analysis of *ntrB, gldH* and *glnA* using q-RT PCR at 3,6,9 and 12 hours in Burk’s medium + 0.2% maltose + 10mM ammonium chloride (BMN) with respect to Burk’s medium + 0.2% maltose. Data are representative of N = 3, n = 3, and are presented as mean ± SD. Analysis was performed using one-way ANOVA. D, E. Growth curve analysis by absorbance of STM WT, STM Δ*ntrB,* STM Δ*rpoE, and* STM Δ*rpoS* under Burk’s medium + 0.2% maltose (BM) condition at various time points in a subculture ratio of 1:1000. Data are representative of N=3, n=3, presented as mean±SD. Statistical analysis was performed using one-way ANOVA on data combined from 3 different experimental sets, 6 hours and 12 hours after the start of the experiment. F, G. Growth curve analysis by absorbance of STM WT, STM Δ*ntrB,* STM Δ*rpoE, and* STM Δ*rpoS* under Burk’s medium + 0.2% maltose + 10mM ammonium chloride (BMN) condition at various time points in subculture ratio of 1:1000. Data are representative of N=4, n=3, presented as mean±SD. Statistical analysis was performed using one-way ANOVA on data combined from 4 different experimental sets, 6 hours and 12 hours after the start of the experiment.

To determine the factors responsible for the growth of STM WT in Burk’s medium supplemented with maltose, we generated STM Δ*ntrB,* STM Δ*rpoS*, and STM Δ*rpoE* bacterial strains. RpoS is required to maintain the CFU of *E.coli* during nitrogen starvation and to promote the bacterial growth on alternative carbon sources (Notley-McRobb *et al*., 2002, Kabir *et al*., 2004). RpoE is an alternative sigma factor that is responsible for maintaining membrane integrity. Because nutrient stress is also linked to membrane stress, we hypothesised that it may be required for survival under nitrogen-deficient conditions. At a subculture ratio of 1:1000, the global nitrogen regulator (STM Δ*ntrB*) deletion showed attenuated growth from 3h to 9h in Burk’s medium with maltose and Burk’s medium with maltose and ammonium chloride compared to STM WT. However, the growth was significantly higher than that of the STM WT at a later time point of 12 hours.

In contrast, STM Δ*rpoS* showed similar growth in Burk’s medium with maltose (BM) and Burk’s medium with maltose and ammonium chloride (BMN) as STM WT **(Figure 3D-E)**. Similar to STM Δ*ntrB,* on a subculture ratio of 1:1000, STM Δ*rpoE* showed reduced growth in Burk’s medium with maltose and Burk’s medium with maltose and ammonium chloride from 3h to 9h compared to STM WT, but the growth was similar to WT at 12 hours **(Figure 3F-G)**. Notably, in Burk’s medium with maltose (BM), STM Δ*rpoE* showed reduced growth compared to STM Δ*ntrB* **(Figure 3E)**, whereas the addition of nitrogen to Burk’s medium with maltose (BMN) led to similar growth of STM Δ*rpoE* and STM Δ*ntrB* at 6 hours **(Figure 3G)**.

At a 1:100 bacterial subculture ratio, STM Δ*ntrB* and STM Δ*rpoE* showed no growth defects in nitrogen-deficient media supplemented with maltose (BM) or maltose, ammonium chloride (BMN), Burk’s media and LB media at a 1:100 ratio of subculture **(supplementary Figure 2A-2D).** At the same time, there was a reduction in the growth of STMΔ*rpoS* in the nutrient-rich LB medium **(supplementary Figure 2D)**.

### 3.3 Growth in nitrogen-deficient conditions is accompanied by reduced pathogenicity

Metabolism is intricately linked to virulence. We analysed whether the growth of *Salmonella* in nitrogen-deficient media affects its pathogenesis. The invasion of STM WT when grown in Burk supplemented with maltose (BM) and Burk supplemented with maltose and ammonium chloride (BMN) was significantly reduced in Caco-2 cells compared to that of LB-grown bacteria **(Figure 4A).** *Salmonella* invasion depends on *Salmonella* pathogenicity island 1 (SPI-1) effectors secreted into host cells, which lead to changes in cytoskeletal arrangement (Hajra *et al*., 2021). q-RT PCR analysis of the SPI-1 genes (*sipB, invF* and *hilA*) showed that STM WT grown in Burk supplemented with maltose (BM) and Burk supplemented with maltose and ammonium chloride (BMN) media exhibited downregulation of these genes compared to those grown in LB*. hilA,* the transcriptional regulator of SPI-1, was downregulated in *Salmonella* grown in Burk medium supplemented with maltose compared to that in Burk medium supplemented with maltose and ammonium chloride **(Figure 4B).** In the gentamicin protection assay for STM WT grown under various conditions, STM WT grown in Burk medium supplemented with maltose (BM) showed the highest proliferation compared to Burk medium supplemented with maltose and ammonium chloride (BMN) media and LB **(Figure 4C)**.

**Figure 4.**
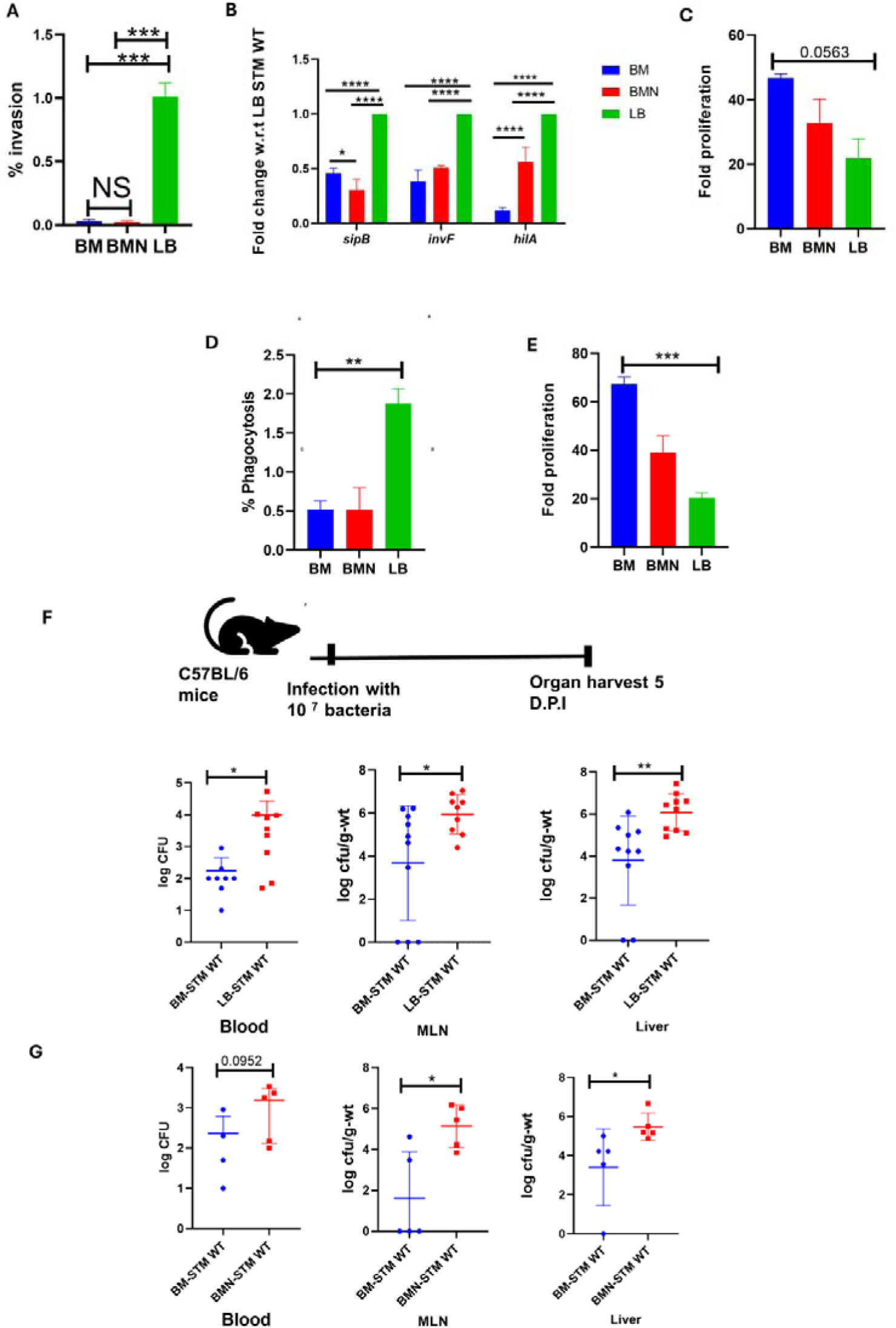
*Salmonella* grown in the nitrogen-deficient condition is attenuated in the pathogenesis. A. Percentage invasion of STM WT grown under different conditions in Caco2 cells - BM (Burk’s medium + 0.2% maltose), BMN (Burk’s medium + 0.2% maltose +10mM ammonium chloride), LB (Luria Bertani medium). Unpaired Student’s t-test was used for analysis. Data are representative of N=4, n=3, and are presented as mean ±SEM. B. mRNA expression of SPI-1 genes (*sipB, invF* and *hilA)* by q-RT PCR in BM, BMN, and LB media grown STM WT. Analysis by two-way ANOVA. Data are representative of N=3, n=3, presented as mean ±SD. C. Fold proliferation of STM WT grown under different conditions - BM (Burk’s medium + 0.2% maltose), BMN (Burk’s medium + 0.2% maltose +10mM ammonium chloride), LB (Luria Bertini medium). An unpaired Student’s t-test was used for analysis. The data are representative of N = 3, n = 3. D. Percentage phagocytosis of STM WT grown under different conditions in RAW 264.7 cells - BM (Burk’s medium + 0.2% maltose), BMN (Burk’s medium + 0.2% maltose +10mM ammonium chloride), LB (Luria Bertani medium). Unpaired Student’s t-test was used for analysis. Data are representative of N=3, n=3, presented as mean ±SD. E. Fold proliferation of STM WT grown under different conditions in RAW 264.7 cells - BM (Burk’s medium + 0.2% maltose), BMN (Burk’s medium + 0.2% maltose +10mM ammonium chloride), LB (Luria Bertini medium). Unpaired Student’s t-test was used for analysis. Data are representative of N=3, n=3, presented as mean ±SD F. Schematic representation of infection for organ burden determination of STM WT grown in the BM and LB. Organ burden in the blood, mesenteric lymph node (MLN), and liver. The Mann-Whitney test was used for analysis, with ten animals per cohort presented as the mean ± SD. G. Schematic representation of infection for organ burden determination of STM WT grown in BM and BMN. Organ burden in the blood, mesenteric lymph node (MLN), and liver. The Mann-Whitney test was used for analysis, with five animals per cohort presented as the mean ± SD.

Macrophages play a crucial role in the dissemination of *Salmonella* to various secondary infection sites. On performing the phagocytosis assay in RAW 264.7, STM WT grown in Burk supplemented with maltose (BM) and Burk supplemented with maltose and ammonium chloride (BMN) showed reduced phagocytosis compared to LB **(Figure 4D)**. Next, when performing the gentamicin protection assay, we observed similar proliferation rates in the Caco-2 cells. STM WT grown in Burk medium supplemented with maltose (BM) showed the highest proliferation compared to STM WT grown in Burk medium supplemented with maltose and ammonium chloride (BMN) media and LB **(Figure 4E)**.

In the *in-vivo* pathogenesis model using C57BL/6 male mice, we observed that *Salmonella* grown in Burk medium supplemented with maltose (BM) was less pathogenic than in the other three conditions: Burk medium (B), Burk medium supplemented with maltose and ammonium chloride (BMN), and LB medium. There was also less weight reduction in mice and more prolonged survival when infected with *Salmonella* grown in Burk medium supplemented with maltose (BM) **(Supplementary Figure 3A, 3B).** Furthermore, the organ burden in C57BL/6 mice infected with Burk supplemented with maltose (BM) grown *Salmonella* was reduced in the blood, mesenteric lymph nodes, and liver compared to LB-grown *Salmonella* five days post-infection **(Figure 4F).** Similarly, there was an increase in organ burden when *Salmonella* was grown in Burk supplemented with maltose and ammonium chloride (BMN) compared to that in Burk supplemented with maltose (BM) **(Figure 4G).** In conclusion, *Salmonella* grown in Burk supplemented with maltose (BM) alone or in combination with ammonium chloride exhibited reduced invasion and phagocytosis in cell lines Caco-2 and RAW 264.7 macrophages; however, there was a higher fold proliferation compared to LB-grown bacteria. In an *in-vivo* model using C57BL/6 mice, we observed a reduced organ burden of *Salmonella* grown in Burk’s medium supplemented with maltose (BM) compared to when supplemented with maltose along with nitrogen (BMN) and LB-grown *Salmonella*.

## 4. Discussion-

Maltose has been studied for its ability to increase host immunity by increasing lysozyme expression in zebrafish against *Vibrio alginolyticus,* decreasing toxin production in *Vibrio Cholerae*, and reducing the adhesion and motility of *Pseudomonas (Lang et al., 1994, Shetye et al., 2014, Jiang et al., 2020).* Recently, we have also demonstrated the role of the maltose metabolism gene *malQ* in reducing *Salmonella* adhesion to epithelial cells (Parmar *et al*., 2025). In this study, we investigated the role of maltose in promoting the growth of *Salmonella* Typhimurium under nitrogen-deficient conditions.

The most common bacterium used to study bacterial survival under nitrogen-deficient conditions is *E.coli*, which, upon nitrogen starvation, stops growing abruptly from maximum to zero in 27 minutes (Bren *et al*., 2013). *E.coli* also had a doubling time of 39 hours during prolonged nitrogen starvation, which was 80-fold higher than the doubling time under nutrient-rich conditions. This growth was attributed to the breakdown of allantoin, which is formed by nucleic acid degradation and the activation of peptidoglycan degradation and transport, serving as a nitrogen donor (Choi *et al*., 2016, Switzer *et al*., 2020). Nitrogen starvation leads to the slowing down of translation and coordination with transcription mediated by ppGpp. Deletion of ppGpp reduces overall protein synthesis and cell viability (Iyer *et al*., 2018). Previous reports in the literature for mimicking enteric bacteria under nitrogen-deficient conditions have used M9 or Gutnick’s minimal media supplemented with 3 mM ammonium chloride as a nitrogen source, which eventually runs out during the experiment, as compared to 10 mM ammonium chloride in a nitrogen-rich condition. Our study found that nitrogen-deficient Burk’s medium (Himedia) contains approximately 4.5 mM nitrogen. Burk’s medium, in literature, has been used to isolate and grow nitrogen fixers (Aasfar *et al*., 2024). Burk’s medium could support the *g*rowth of *Salmonella* under nitrogen-deficient conditions by adding maltose to both liquid and solid media (**Figure 1A-B).** Deletion of the maltose metabolism gene (*malQ)* or the maltose transporter (*malEFG)* significantly attenuated growth (**Figure 1C-D).**

Bacterial length has been shown to be dependent on the carbon status of bacteria, where growth is dependent on the levels of UDP-glucose in *E.coli (Weart et al., 2007)*. STM grown in Burk’s medium had reduced length compared to that grown in Burk’s media supplemented with maltose alone or maltose with ammonium chloride and LB medium (**Figure 2A-B**). These results suggest that the carbon source was limited in Burk’s media. Burk’s medium contains sucrose as a carbon source, which *Salmonella* generally cannot utilise. However, some serovars, such as *Salmonella* Mbandaka and *Salmonella* Bareilly, can ferment sucrose (Reid *et al*., 1993, Jose, 2020). Recent reports have shown that certain *Salmonella* Typhimurium types can also utilise sucrose due to the acquisition of genes important for metabolism (Stevens *et al*., 2024). *Salmonella* grown in Burk medium supplemented with maltose (BM) condition had high alpha-ketoglutarate levels, and growth was found to be significantly attenuated upon deletion of the NtrBC two-component system and sigma factor *rpoE* (**Figure 3D-G).** STM Δ*rpoE* showed significantly attenuated growth compared with STM Δ*ntrB* in Burk’s medium supplemented with maltose. STM Δ*rpoE* behaved similarly to STM Δ*ntrB* in Burk’s medium supplemented with maltose and nitrogen. This suggests that STM Δ*rpoE* is more sensitive to nitrogen limitation than STM Δ*ntrB.* Growth reduction upon deletion of *ntrB* is consistent with a previous report by Schumacher et al, where they showed similar results in *E.coli* that *ntrB* is required for bacterial growth under both nitrogen-rich and nitrogen-deficient conditions(Schumacher *et al*., 2013). However, the growth advantage of STM Δ*ntrB* at at a later time point of 12 hours in both Burk supplemented with maltose (BM) and Burk supplemented with maltose and nitrogen (BMN) indicated that adavantge is independent of nitrogen supplementation. Future studies in this direction are needed to address how STM Δ*ntrB* on maltose supplementation with limation of nitrogen is leading to the growth advantage.

The starvation of various bacterial pathogens is known to influence virulence. The fish pathogen *Flavobacterium columnare*, which had been starved in a lake for 5 months, exhibited greater virulence compared to its ancestral strain (Sundberg *et al*., 2014). Phosphate limitation in *Bacillus anthracis* increases internalisation in RAW 264.7 macrophages and its pathogenesis in mice (Aggarwal *et al*., 2015). Carbohydrates (sorbitol, galactose, and maltose) also influence biofilm formation and siderophore production in *Klebsiella (Almeida et al., 2025).* In our study, we found that bacteria grown under nitrogen-deficient conditions, when supplemented with maltose alone or in combination with ammonium chloride, were defective in invasion and phagocytosis in cell line models (Caco-2 and RAW macrophages) compared to bacteria grown in LB medium. The reduced invasion of Caco-2 cells was found to be due to the decreased expression of the T3SS encoded by SPI-1. We have not explored the reasons for reduced phagocytosis by macrophages; it is largely presumed to be a passive uptake process, but recent reports suggest that T3SS-dependent uptake occurs in macrophages at the initial time points (Di Martino *et al*., 2019). In contrast to *in vitro* cell line results, where we observed no difference in the pathogenesis of BM and BMN, nitrogen supplementation with maltose in Burk’s medium (BMN) increases the organ burden in the C57BL/6 mice, compared to bacteria grown in Burk’s medium supplemented with maltose (BM) **(Figure 4).**

## Ethics statement

All the animal experiments were approved by the Institutional Animal Ethics Committee (IAEC) of the Indian Institute of Science, Bangalore. The ethical clearance number for this study was CAF/ethics/852/2021. All the animals were acquired from the central animal facility of the Indian Institute of Science, Bangalore.

## Availability of data and materials

The data supporting the findings of this study are available from the corresponding authors upon reasonable request.

## Author contribution

**Kirti Parmar**- writing-review and editing, visualisation, experimental design, project administration, investigation and data analysis.**Yogyta Kumari**-investigation, **Kapudeep Karmakar-** manuscript editing and visualisation. **Muralidhar Nayak**-alpha-ketoglutarate estimations. **Sumanta Bagchi**- Elemental analysis. **Dipshikha Chakravortty**- writing-review and editing, visualisation, experimental design, investigation, data analysis, resources and supervision.

## Acknowledgements

Abhilash Vijay Nair is acknowledged for helping with the oral gavage of the animals. We acknowledge the support of the department’s real-time facility and central animal facility, as well as the Advanced Facility for Microscopy and Microanalysis at IISc, for the experiments. Professor Sumanta Bagchi acknowledges the DST (FIST) for the elemental analysis instrument. We sincerely thank Amit Sahu and Niladri for their assistance with the growth curve machine (Bioscreen).

## Supplementary information

**Supplementary Figure 1.**
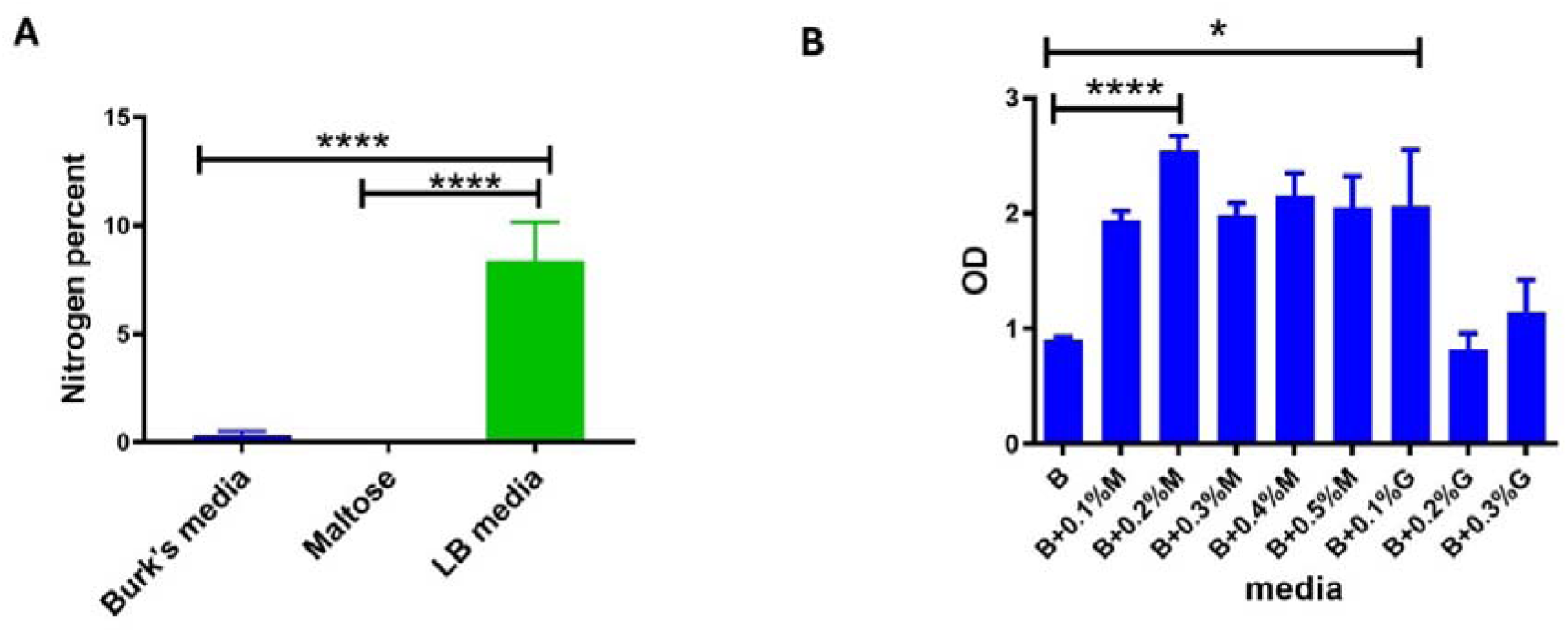
*Salmonella* growth under various conditions following a 1:100 subculture. A. CHN analysis was used to determine the nitrogen percentage in Burk’s medium, Maltose and LB medium. An unpaired Student’s t-test was used for the analysis. Data represent n=6, presented as mean ± SD. B. *Salmonella* Typhimurium growth in Burk’s medium with supplementation of glucose and maltose at various concentrations. Unpaired Student’s t-test was used for analysis. Data are representative of N=3, n=3, presented as mean ±SD.

**Supplementary Figure 2.**
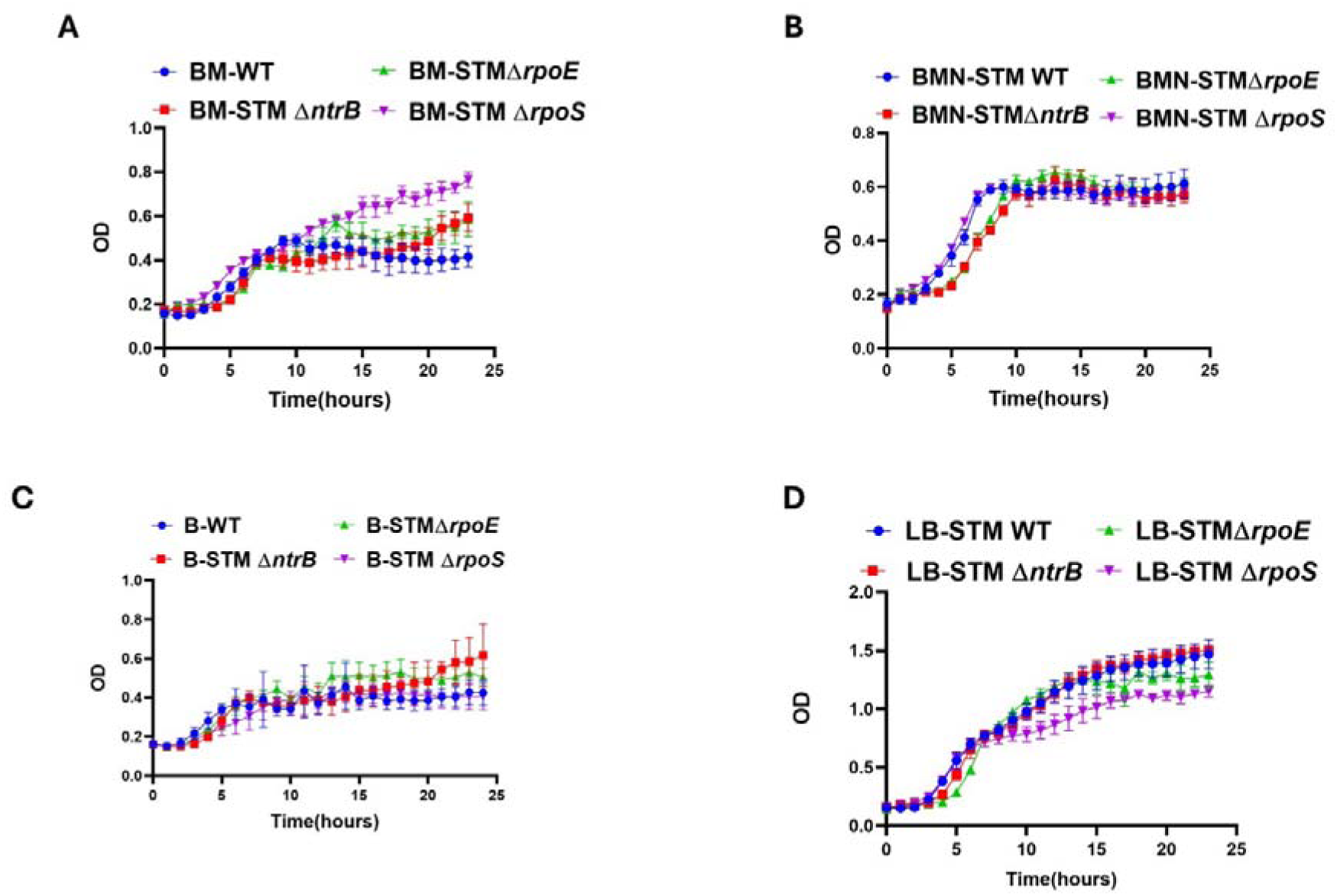
Growth analysis of various bacterial strains at a 1:100 subculture ratio. A-D. Growth curve analysis by absorbance of STM WT, STM Δ*ntrB,* STM Δ*rpoE,* and STM Δ*rpoS* in Burk’s medium with maltose (BM), Burk’s medium with maltose and ammonium chloride (BMN), Burk’s medium (B) and LB medium. Data are representative of N=3, n=5, presented as mean ± SD.

**Supplementary figure 3.**
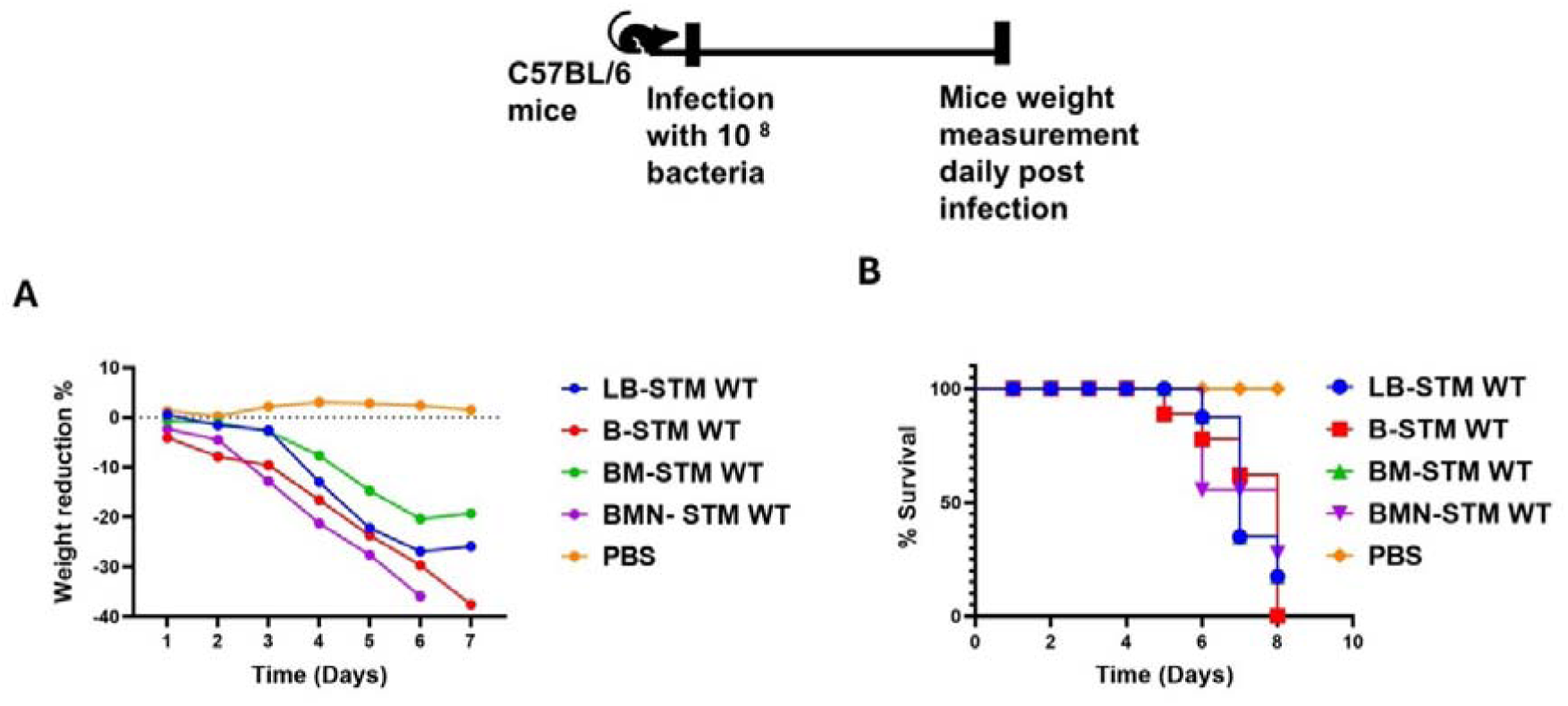
Survival of mice when infected with STM grown in different conditions. A and B. Weight reduction percentage and survival in C57BL/6 mice post-infection by STM WT grown in B (Burk’s medium), BM (in Burk’s medium with maltose), BMN (in Burk’s medium with maltose and ammonium chloride) and LB media. Five animals per cohort were used for the survival curve.

**Supplementary Table 1.**
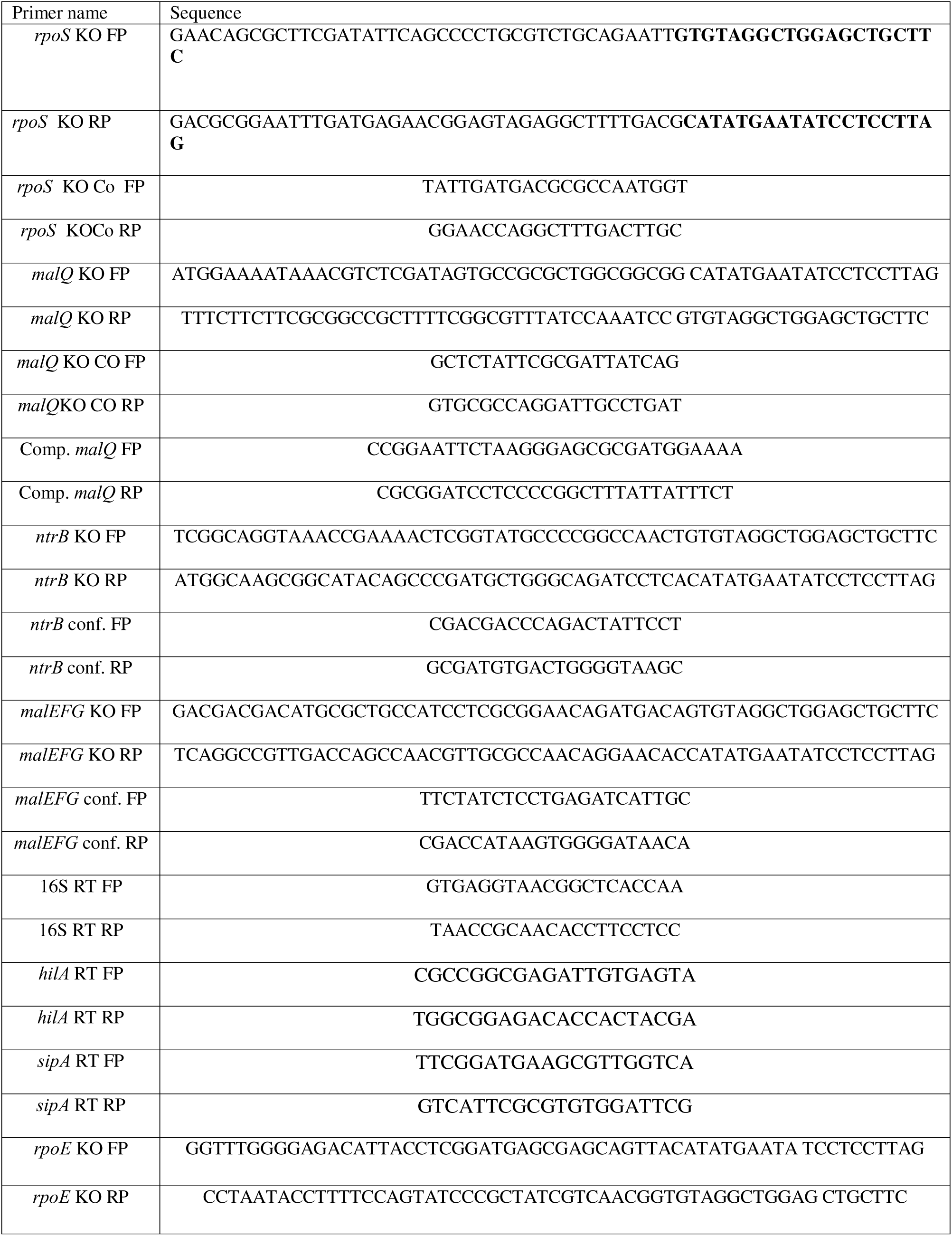
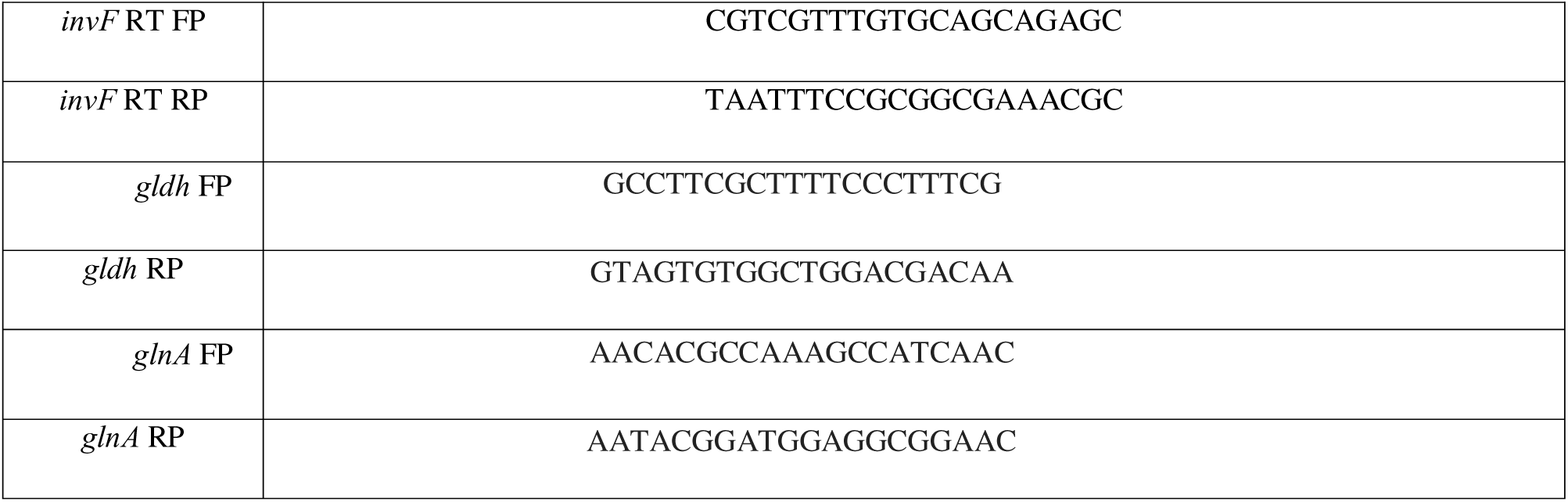
List of primers used in the study.

